# Seasonal transcriptional profiling of malaria parasites using a new single-cell atlas of a Malian *Plasmodium falciparum* isolate

**DOI:** 10.1101/2025.04.14.648697

**Authors:** Lasse Votborg-Novél, Martin Kampmann, Manuela Carrasquilla, Georgia Angeli, Christina Ntalla, Hamidou Cisse, Yuhana Sogoba, Gabriela M Guerra, Safiatou Doumbo, Didier Doumtabe, Silke Bandermann, Yara Cuesta Valero, Kassoum Kayentao, Mir-Farzin Mashreghi, Moussa Niangaly, Aissata Ongoiba, Peter D Crompton, Thomas D Otto, Boubacar Traoré, Silvia Portugal

## Abstract

Asymptomatic persistence is crucial for the survival of malaria parasites in seasonal transmission settings, where transmission halts during dry months, while mosquitos are scarce and new infections rare. How *Plasmodium falciparum* avoids causing host symptoms or being cleared is not fully understood. Parasites that persist through the dry season circulate longer within the ∼48h replication cycle compared to parasites isolated from clinical cases, where only early stages circulate. The mechanisms promoting increased circulation and avirulent infections bridging two wet seasons remain unclear. We generated a single-cell RNA-seq reference atlas with *P. falciparum* erythrocytic stages of a recently adapted isolate, and with it defined the developmental stage of over 25000 parasites from six asymptomatic children at the dry-wet season transition, and nine children with clinical malaria in the wet season. Our data confirms that older parasites circulate in the dry season, and reveals transcriptional modulation of exported proteins that affect cytoadhesion of infected erythrocytes, which we validated by immuno-fluorescence microscopy.

## Introduction

The mosquito-borne *Plasmodium falciparum* parasite remains a leading cause of morbidity and mortality in low-income countries, causing over 200 million clinical malaria cases and over half a million deaths of African children each year (1). Malaria symptoms, typically a mild fever that can progress to severe disease and death in young children (2), are caused by the blood infection of parasites going through rounds of ∼48h asexual replication cycles inside red blood cells (RBCs). Within each cycle, RBCs are progressively remodelled and parasites express variant antigens on protrusions of the erythrocyte surface called knobs (3), which allow adhesion of infected RBCs (iRBCs) to the vascular endothelium, avoiding splenic clearance (4), and promoting rapid expansion of parasitaemia. Malaria transmission varies geographically from unstable to holoendemic (5), occurring year-round in areas with continuous mosquito presence; or seasonally, dropping from high to nearly zero during dry periods (6), due to the absence of mosquito vectors that depend on water for reproduction. In vast regions of Africa where rain seasonality limits vector availability for months, malaria cases are restricted to the wet season, while clinically silent *P. falciparum* infections can persist through the dry season and are important reservoirs for transmission (7–10).

In Mali, transmission is sharply seasonal, with the yearly resurgence of *P. falciparum* transmission every wet season ensured by the asymptomatic reservoir persisting undetected during the ∼6-month dry season in 10–30% of children (7, 10, 11). The molecular mechanisms that allow the parasite to evade immune clearance while maintaining low parasitaemias that do not risk host survival, and hence potentiate future transmission, are unknown. We have shown previously that parasites isolated in the dry season are transcriptionally distinct from those causing febrile malaria in the wet season, and that the transcriptional differences reflect longer circulation within each ∼48h intra-erythrocytic cycle of iRBCs at the end of dry season (7), whereas in febrile disease mostly early stages within the 48h cycle are found in circulation, while later parasite forms cytoadhere (12). Bulk RNA-seq analysis of *P. falciparum* from the dry and the wet seasons was heavily affected by differences in developmental stage of circulating parasites in the two conditions (7), and therefore could not resolve how dry season parasites deviate from those in clinical malaria cases.

Single cell transcriptomics is critical to compare pools of parasites with equivalent expression levels of stage defining transcripts, to allow the discovery of differentially expressed genes (DEGs) truly linked to subclinical persistence. Single-cell RNA-seq of laboratory-adapted parasite lines has revealed novel transcriptional signatures of *P. falciparum* in vitro (13), and of *P. vivax* infecting simian experimental models (14), but single-cell transcription data from natural infections is scarce. Recently, a single-cell malaria atlas of *P. falciparum* blood stage development was built from ∼37000 iRBCs of two well established *P. falciparum* lines, NF54 and 7G8, and integrated with ∼8000 single iRBCs from four naturally infected individuals in Mali (15). Here, we used *P. falciparum* of a recently adapted line to generate a single-cell atlas including all intra-erythrocytic asexual stages and the sexual development axis of the parasite in the blood. This atlas was used to define the developmental age, based on single-cell transcriptomes, of individual parasites collected in Mali from persisting dry season infections and those of clinical malaria cases in the ensuing wet season, which we then used to investigate DEGs between dry and wet season parasites.

## Results

### Single-cell RNA-seq atlas of asexual and sexual stages of a *P. falciparum* isolate

We profiled single-cell transcriptomes of the blood-stages of *Pf*M2K1, a Malian isolate of *P. falciparum* collected in 2018 that was cloned and adapted to in vitro culture. Cells covering all intra-erythrocytic developmental stages (Supplementary Table 1) were filtered to exclude cells with <200 genes, >20,000 unique molecular identifiers (UMIs), and >5% mitochondrial UMIs, and were integrated and clustered according to their parasite developmental stage (16, 17). Initial unsupervised clustering yielded six clusters (Supplementary Fig. 1a), with one low-quality iRBC cluster removed after further filtering (Supplementary Fig. 1b). Post quality control, 26,447 M2K1 iRBCs were integrated and visualized on UMAP (Uniform Manifold Approximation and Projection) (Fig. 1a), displaying a circular arrangement with branching cells corresponding to asexual replication and gametocyte development, consistent with other malaria single-cell atlases (15, 17).

**Figure 1.**
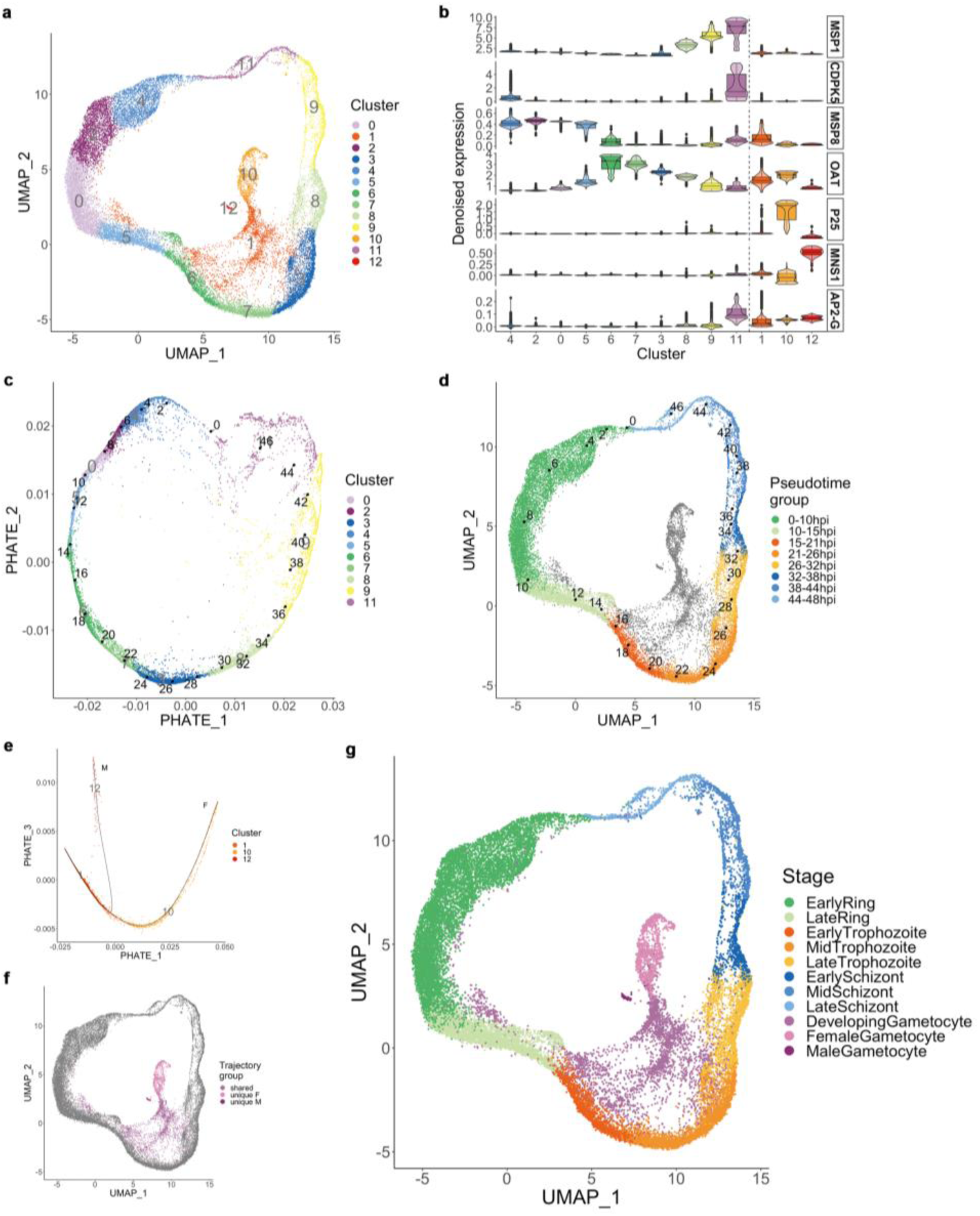
Single-cell RNA-seq atlas of asexual and sexual stages of a *P. falciparum* isolate. **a,** UMAP of 26447 single *Pf*M2K1 iRBCs coloured by 13 Smart Local Moving (SLM) clusters. **b,** Expression of parasite developmental stage marker genes denoised using MAGIC. **c,** PHATE of the 22282 asexual iRBCs across 10 clusters annotated by hours post invasion (hpi), and **d,** UMAP with the asexual iRBCs coloured and annotated by hpi. **e,** PHATE of the 4165 sexual iRBCs across 3 clusters annotated by two inferred trajectories, and **f,** UMAP with the asexual iRBCs annotated according to the inferred trajectories. **g,** UMAP of the 26447 single-iRBCs coloured by annotated parasite developmental stages.

Quality-filtered iRBCs were grouped into 13 unsupervised clusters (0–12, decreasing in cell count) (Fig. 1a). Gene expression was denoised using Markov Affinity-based Graph Imputation of Cells (MAGIC) (18) to address dropout across similar cells. Stage-specific genes like schizogony-marker *msp1* (PF3D7_0930300) (19); egress-activator *cdpk5* (PF3D7_1337800) (20)); reinvasion-marker *msp8* (PF3D7_0502400) (21); and trophozoite-expressed *oat* (PF3D7_0608800) (22, 23), established a counter-clockwise asexual cycle orientation. RBC reinvasion occurred in cluster 4, progressing through clusters 2, 0, 5, 6, 7, 3, 8, 9, and 11, with trophozoites in cluster 6, and schizonts in cluster 8 (Fig. 1b).

Gametocyte markers *p25* (PF3D7_1031000) and *mns1* (PF3D7_1311100) (24) identified female and male gametocytes in cluster 10 and 12, respectively (Fig. 1b), supported by additional sex-specific marker genes (24) (Supplementary Fig. 1c and d). Cluster 1 appeared alongside clusters following reinvasion (4, 2, 0, and 5), branching off from the subsequent clusters (6, 7, 3, and 8) and transitioning into the gametocyte clusters (10 and 12) (Fig 1a). This aligned with cluster 1 representing sexually developing iRBCs, evidenced by elevated *ap2g* (PF3D7_1222600) expression (Fig. 1b), a regulator of sexual commitment (25); and by early sexual development markers (26) (Supplementary Fig. 1e).

To represent the global structure of the projected iRBCs, we used Potential of Heat diffusion for Affinity-based Transition Embedding (PHATE), that emphasizes local transitions (27), and enabled pseudotime-ordering of iRBCs along the ∼48h asexual cycle. The resulting structure resembled UMAP (Supplementary Fig. 2a and b), with minor differences in cluster proportions and spacing. PHATE confirmed cluster 1 as sexually developing iRBCs but did not differentiate female and male gametocytes (Supplementary Fig. 2a and b). Using Slingshot (28) on PHATE-based k-means clustering, we identified one lineage following the asexual cycle, and another the sexual lines ending at the gametocyte clusters 10 and 12 (Supplementary Fig. 2c, d). However, Slingshot placement of asexual clusters (Supplementary Fig. 2c) conflicted with *cdpk5* and *msp8* markers pinpointing the reinvasion point (Fig. 1b); and Slingshot failed to separate developing gametocytes from the asexual lineage (Supplementary Fig. 2c, d). Thus, we analyzed asexual and sexual clusters separately.

Projection of the 10 asexual clusters into PHATE revealed a circular arrangement, with the trajectory start aligning with egress and invasion marker genes, enabling pseudotime scaling to the 48h cycle to define hours post-invasion (hpi) for each iRBC (Fig. 1c). Asexual iRBCs were grouped into stages: early ring (0–10hpi, n = 9099), late ring (10–15hpi, n = 3346), early trophozoite (15–21hpi, n = 1694), mid trophozoite (21–26hpi, n = 3309), late trophozoite (26–32hpi, n = 1863), early schizont (32–38hpi, n = 958), mid schizont (38–44hpi, n = 1198), and late schizont (44–48hpi, n = 815) (Fig. 1d). The three sexual clusters showed clear separation of male (cluster 12) and female (cluster 10) gametocytes in PHATE, with two trajectories ending in each cluster (Fig. 1e). This included 1214 iRBCs unique to female (n = 1133) or male (n = 81) trajectories, and 2951 iRBCs shared as developing gametocytes (Fig. 1e, f). A final annotated atlas combined asexual and sexual cluster annotations (Fig. 1g).

M2K1 iRBCs were grouped into 48 pseudotime bins of asexual iRBCs (1–48, n = 38–1845), six of developing gametocytes (DG1–DG6, n = 84–919), two of female gametocytes (FG1 and FG2, n = 478 and 655), and one of male gametocytes (MG, n = 81). Pseudo-bulk samples were created by combining iRBC expression within each bin. Using 48h bulk data by Bozdech & Llinas (29) we determined developmental age by maximum likelihood estimation (MLE) (30), revealing a strong correlation with asexual progression (Pearson’s r: 0.9947, P < 0.00015) (Supplementary Fig. 3a), validating the reinvasion point. For sexual stages, pseudo-bulk samples were compared to 16-day bulk data by van Biljon et al. (31), grouped into asexual or sexually committed, and early, mid, and late gametocyte stages. Median correlations for pre-induction/commitment were stable, while early, mid, and late gametocyte groups showed increasing correlation across DG1–DG6 and toward female/male gametocytes (Supplementary Fig. 3b), consistent with sexual development trajectories.

We compared the M2K1 atlas with single-cell malaria atlases from Howick & Russel et al. (17) and Dogga, Rop & Cudini et al. (15), using *scmap* (32). Projecting the Howick atlas onto M2K1 atlas revealed minimal ring-stage overlap (1.0%) (Supplementary Fig. 3c), indicating limited ring coverage in the Howick atlas. Pseudotime correlation between M2K1 and Howick atlases was moderate (Pearson’s r: 0.745, P < 0.0001) (Supplementary Fig. 3d), with discrepancies in pseudotime ordering and reinvasion point, supported by MLE analysis (Pearson’s r: 0.514, P < 0.0002) (Supplementary Fig. 3e). Projecting the Dogga atlas to M2K1 showed good asexual and gametocyte coverage but misaligned early rings with M2K1’s reinvasion point (Supplementary Fig. 3f). M2K1 projection to the Dogga atlas revealed overall alignment with gradual asexual stage transitions (Supplementary Fig. 3g), likely due to differing pseudotime methods (PHATE vs. UMAP). Half of M2K1’s late schizonts projected to Dogga’s early rings, highlighting reinvasion point differences. Match of M2K1’s developing gametocytes to Dogga’s gametocytes increased along the sexual trajectories, (3.3% in DG1 to 95.9% in DG6) (Supplementary Fig. 3g), indicating M2K1’s broader detection of early developing gametocytes. Altogether, we built a single-cell atlas of a recently adapted *P. falciparum* clone, encompassing all asexual and sexual blood stages. Using trajectory and pseudotime inference guided by unsupervised clustering and marker gene expression, we annotated 26,447 iRBCs. We demonstrated higher coverage of ring stages compared to the existing atlas of *P. berghei* (17), more precise determination of the merozoite reinvasion point, better pseudotime-ordering of cells, and better coverage of developing gametocytes compared to the Dogga *P. falciparum* atlas (15).

### Single-cell transcriptomics of *P. falciparum* natural infections

We profiled single-cell transcriptomes of *P. falciparum* iRBCs of six asymptomatic individuals at the dry-to-wet season transition (endDry), and of nine clinical malaria cases diagnosed in the ensuing wet season (MAL), in Kalifabougou, Mali. Participants, aged 4–17, were part of a previously described cohort (33). Median parasitaemia was lower at the end of the dry season (0.071%, IQR: 0.051–0.141) than in malaria cases in the wet season (2.790%, IQR: 0.370–3.240) (Wilcoxon rank sum test, P=0.004795). Before single-cell capture, iRBCs were enriched by flow cytometric sorting (Supplementary Fig. 4), achieving 74,4% (IQR: 60.8–90.4) and 89.8% (IQR: 76.5–91.4) of iRBCs in the dry-to-wet season transition and clinical malaria, respectively (Table 1).

**Table 1.** Overview of *P. falciparum* natural infections of asymptomatic carriers at the dry to wet season (endDry, n = 6) and symptomatic clinical malaria cases in the wet season (MAL, n = 9).

| Sample | Individual age [years] | Individual gender | Collection date | Individual temperature [°C] | Parasitemia pre sort | Sorting time [min] | Parasitemia post sort | iRBCs post QC |
| --- | --- | --- | --- | --- | --- | --- | --- | --- |
| endDry20K281 | 17 | Male | Jul 2020 | 35,9 | 0,160 % | 90 | 96,1 % | 2012 |
| endDry20K334 | 17 | Male | Jul 2020 | 36,2 | 0,060 % | 110 | 53,2 % | 236 |
| endDry20K405 | 16 | Female | Jul 2020 | 35,8 | 0,048 % | 115 | 59,4 % | 295 |
| endDry20K554 | 12 | Male | Jul 2020 | 36,8 | 0,360 % | 90 | 92,6 % | 408 |
| endDry20K616 | 10 | Male | Jul 2020 | 35,9 | 0,082 % | 120 | 83,7 % | 683 |
| endDry20K998 | 4 | Male | Jul 2020 | 36,4 | 0,034 % | 145 | 65,1 % | 54 |
| MAL18K535 | 12 | Male | Sep 2018 | 39,2 | 0,370 % | 19 | 72,0 % | 955 |
| MAL18K562 | 10 | Female | Sep 2018 | 38,6 | 4,980 % | 10 | 95,5 % | 450 |
| MAL18K623 | 8 | Male | Oct 2018 | 39,0 | 6,250 % | 10 | 94,5 % | 1750 |
| MAL19K557 | 11 | Female | Aug 2019 | 37,7 | 3,200 % | 15 | 91,4 % | 2156 |
| MAL19K891 | 4 | Male | Aug 2019 | 38,1 | 2,790 % | 20 | 75,2 % | 11256 |
| MAL20K269 | 17 | Male | Oct 2020 | 37,7 | 0,170 % | 65 | 90,4 % | 260 |
| MAL20K380 | 16 | Male | Sep 2020 | 38,1 | 0,530 % | 60 | 84,0 % | 277 |
| MAL20K435 | 16 | Male | Oct 2020 | 37,7 | 0,075 % | 60 | 76,5 % | 56 |
| MAL20K557 | 12 | Female | Oct 2020 | 38,5 | 3,240 % | 85 | 89,8 % | 6047 |

After enrichment, single-cell capture and sequencing were performed, including a globin transcript depletion step (34). Sequenced iRBCs were filtered to exclude cells with <100 genes, >20,000 UMIs, >5% mitochondrial UMIs, or <95% *P. falciparum* UMIs (Supplementary Fig. 5a). We retained 3688 cells from the dry to wet season transition samples, and 23207 cells of the wet season clinical malaria cases (Table 1 and Supplementary Fig. 5b and c). Developmental stages were determined by projecting cells onto the M2K1 atlas (Fig. 1g). Both dry season and clinical malaria iRBCs spanned similar hpi ranges (4.4–27.1 vs. 3.9–24.3) (Fig. 2a), but dry season parasites were more advanced in the 48h cycle (Fig. 2a), aligning with prior bulk analysis (7, 35). Dry season samples had fewer early rings (22.42% vs. 95.66%) but more late rings (68.14% vs. 3.92%) and trophozoites (8.30% vs. 2.04%) than clinical malaria cases. No schizonts or male gametocytes were found; and female gametocytes were detected only in dry season cells (0.14%). Median hpi was higher in dry season parasites (11.0 hpi, IQR: 10.9–11.3) than in clinical cases (8.7 hpi, IQR: 8.7–8.8, Wilcoxon rank sum test, P=0.02557) (Fig. 2b). Projections to Howick (17) and Dogga (15) atlases confirmed more developed parasite stages in the dry season infections than clinical malaria cases (Fig. 2c), but with lower cosine similarity and certainty compared to the M2K1 atlas (Wilcoxon signed rank test, P<0.0001) (Fig. 2d). Projection certainty was higher for clinical malaria iRBCs than dry season cells across all atlases.

**Figure 2.**
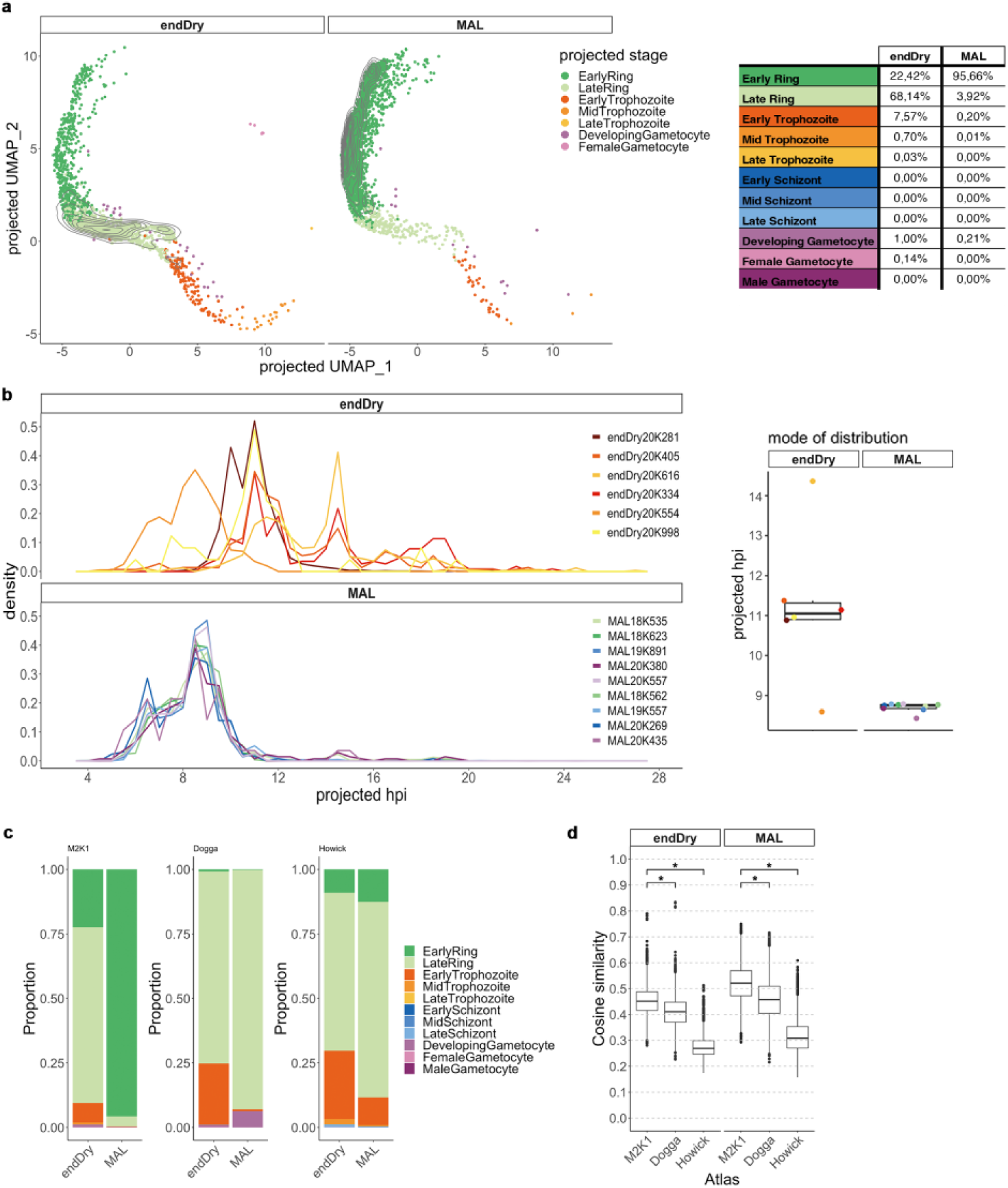
Projection of natural infections’ iRBCs to malaria cell atlases. **a,** UMAP of the projection to the M2K1 atlas of 3688 single-iRBCs from six asymptomatic infections at the transition from dry to wet season (endDry) and 23207 single-iRBCs from nine clinical malaria cases (MAL) in the wet season (left), and proportion of parasite developmental stages in natural infections of endDry and MAL (right) **b,** Distribution of asexual iRBCs from individual samples along the projected hpi (left), and mode of distributions of projected hpi (right). **c,** Parasite developmental stage composition in endDry and MAL based on projection to the M2K1 atlas, and to atlases of *P. falciparum* (Dogga) and of *P. berghei* (Howick). **d,** Cosine similarity measure of the projections, * P < 0.0001, Wilcoxon signed rank test.

### Differential expression of stage-matched iRBCs of natural infections

To compare similar developmental stages, iRBCs from dry season and clinical malaria cases were grouped into 0.25 hpi bins based on M2K1 atlas pseudotime projections (Fig. 3a). Bins were merged to ensure >100 stage-matched cells per condition, resulting in 10 bins between 6 and 11.25 hpi (Fig. 3b, c, Supplementary Table 2). Lower hpi bins were limited by dry season cell counts, while higher bins were constrained by clinical malaria cell numbers (Fig. 3b). Within each bin, hpi was comparable between dry season and clinical malaria iRBCs (Fig. 3c). DEG analysis identified 26 DEGs (fold-change >1.5, adj. P<0.05) across 552–719 genes per bin, with a total of 889 genes tested (Supplementary Table 3). Fourteen genes were up-regulated in the dry season iRBCs, of which 11 were DEGs in two or more consecutive hpi bins. Six of the 12 down-regulated DEGs in the dry season iRBCs, were so in two or more consecutive hpi bins (Fig. 3c). Of the 26 DEGs, 23 were previously tested in bulk data (7), with 14 showing consistent regulation (7 up, 7 down) (Supplementary Table 4). Additionally, 245 genes differentially expressed in bulk data (7), were not DEGs in our single-cell analysis, confirming effective control for expression differences driven by developmental age. Gene ontology (GO) analysis revealed overrepresentation of terms related to cytoadhesion, host cell remodelling, and protein export, while terms linked to translation and ribosome pathway appeared suppressed (gene set enrichment analysis, adj. P<0.05) (Fig. 3d, Supplementary Table 5).

**Figure 3.**
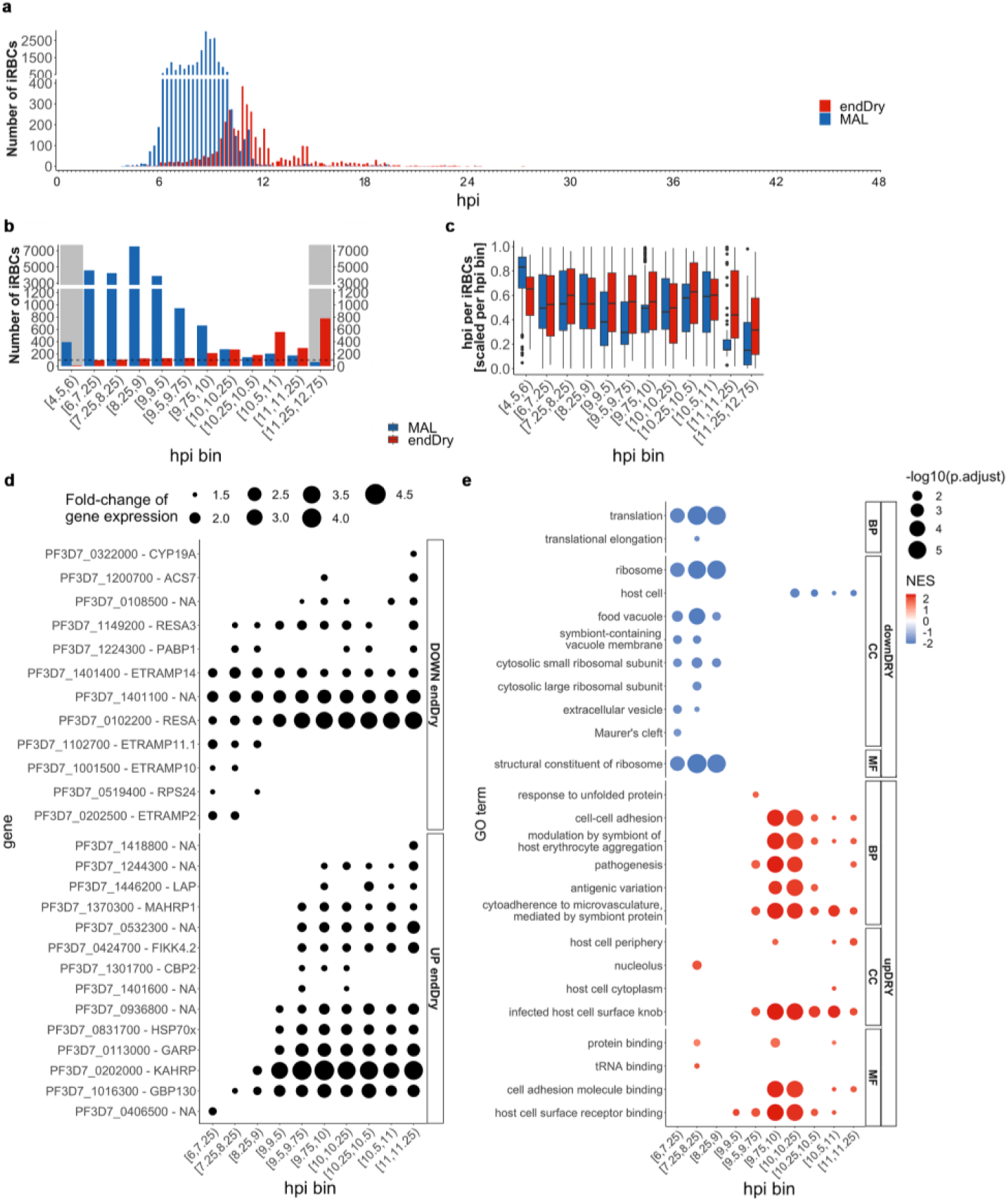
Differential expression of stage-matched iRBCs of natural infections. **a,** Histogram of 3688 iRBCs of asymptomatic infections at the transition from dry to wet season (endDry) and of 23,207 iRBCs of clinical malaria cases in the wet season (MAL) according to hpi (bin size = 0.25), and **b,** hpi bins of at least 100 stage-matched iRBCs (dashed line). Grey shading indicates excluded hpi bins. **c,** Relative hpi of endDry and MAL iRBCs within hpi bins. **d,** DEGs of endDry vs MAL within each hpi bin. Dot sizes indicate fold-change. **e,** Significantly-enriched gene ontology terms within each hpi bin (adjusted P<0.05, gene set enrichment analysis).

### Dry season circulating iRBCs express high KAHRP at the RBC surface

To validate DEG findings, we assessed protein levels of dry season upregulated DEG *kahrp* and *mahrp* via immunofluorescence assay (IFA) on thin blood smears from six asymptomatic individuals at the dry season’s end (age 9-16), and six clinical malaria cases (age 9-14). Samples were stained with anti-KAHRP (36) and anti-MAHRP1 (37, 38) antibodies, anti-EXP2 to highlight the parasitophorous vacuole (39, 40), and Hoechst for counterstaining of nuclei. We analysed 246 iRBCs (endDry: 60, MAL: 186), measuring nuclei size, MAHRP1 staining area, and KAHRP membrane localization.

Consistent with more developed parasites in dry season infections, nuclei were larger in dry season iRBCs (2.15 µm², IQR: 1.63–2.57) than in clinical malaria cases (1.32 µm², IQR: 1.16–1.67) (Fig. 4a). Cells were grouped by nuclei size into quartiles, MAHRP1 stained area was greater in dry season parasites with comparable nuclei size (Fig. 4b). KAHRP positivity was higher in dry season iRBCs across all nuclei sizes (Fig. 4c), with clearer membrane localization in dry season cells compared to clinical cases (Fig. 4d, e).

**Figure 4.**
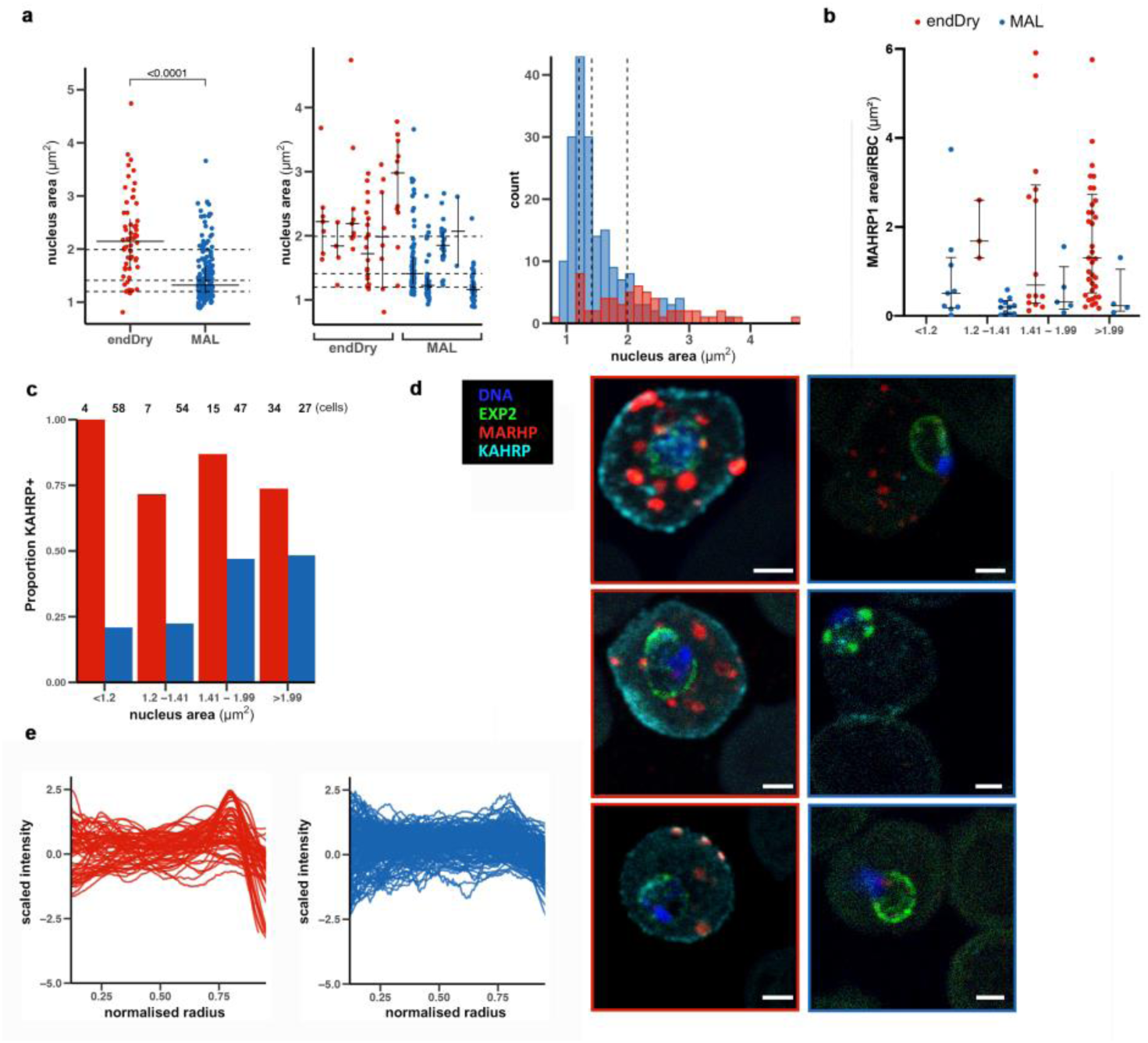
Dry season circulating iRBCs express high KAHRP at the RBC surface. **a,** *P. falciparum* nucleus area measured from thin smears in subclinical infections at the end of the dry season (endDry, n=60) and in clinical malaria cases (MAL n=186) in the wet season (left all cells, centre all cells by individual donor, right histogram of all cells with dashed lines indicating quantiles). Data indicate median ± IQR; two-tailed Mann– Whitney test, *P* < 0.0001. **b,** MAHRP1 area of *P. falciparum* parasites endDry (n=39) and MAL (blue, n=22) grouped by nuclei size. **c,** proportion of KAHRP+ iRBCs different nuclei size. **d,** Representative confocal images of thin blood films of endDry (red) and MAL (blue), stained with anti-EXP2, anti-MARHP1, anti-KAHRP and Hoechst. Scale bar, 2 μm. **e,** Anti-KAHRP radial profiles of all iRBCs of endDry (n=60) and clinical malaria cases (n=186).

### Simulating dry and wet conditions experimentally

To explore mechanisms behind gene expression differences, we simulated dry season vs. clinical malaria conditions in vitro by examining the effects of cytoadhesion and febrile temperatures on *P. falciparum* gene expression. We enriched cultured M2K1 and FCR3 parasites for knobby, adherent iRBCs (panned) (41) using TNF-stimulated human dermal microvascular endothelial cells (HDMECs) (42) and compared them to non-enriched (unpanned) controls. Additionally, we mimicked malarial fever by culturing FCR3 at 41°C for 8h over three cycles (febrile) (43) versus continuous 37°C (afebrile). After single-cell capture, sequencing, and quality filtering (∼2500–6900 cells per condition), we projected cells onto the M2K1 atlas, confirming stage composition by microscopy (Fig. 5a). DEG analysis between 6–11.25 hpi iRBCs revealed no significant DEGs in FCR3 unpanned vs. panned comparisons, (Supplementary Table 6), but 32 DEGs in M2K1 unpanned vs. panned cells, with low consistency across hpi groups (Fig. 5b, Supplementary Table 7). Five DEGs overlapped with the endDry vs. MAL list, but regulation direction was inconsistent (Supplementary table 8); e.g., *garp*, *kahrp*, *hsp70x*, and *mahrp1* were up-regulated in dry season iRBCs but down-regulated in unpanned iRBCs, aligning with their roles in cytoadhesion but not explaining longer circulation of dry season iRBCs. In febrile vs. afebrile FCR3 comparisons, 61 DEGs were identified (Supplementary Table 9), with 6 overlapping the endDry vs. MAL gene list. *garp*, *hsp70x*, and *lap* were up-regulated in both comparisons, while *kahrp*, *fikk4.2*, and *cyp19a* showed opposite regulation (Supplementary Table 8), suggesting febrile temperatures may partially drive expression differences between dry season and clinical malaria infections.

**Figure 5.**
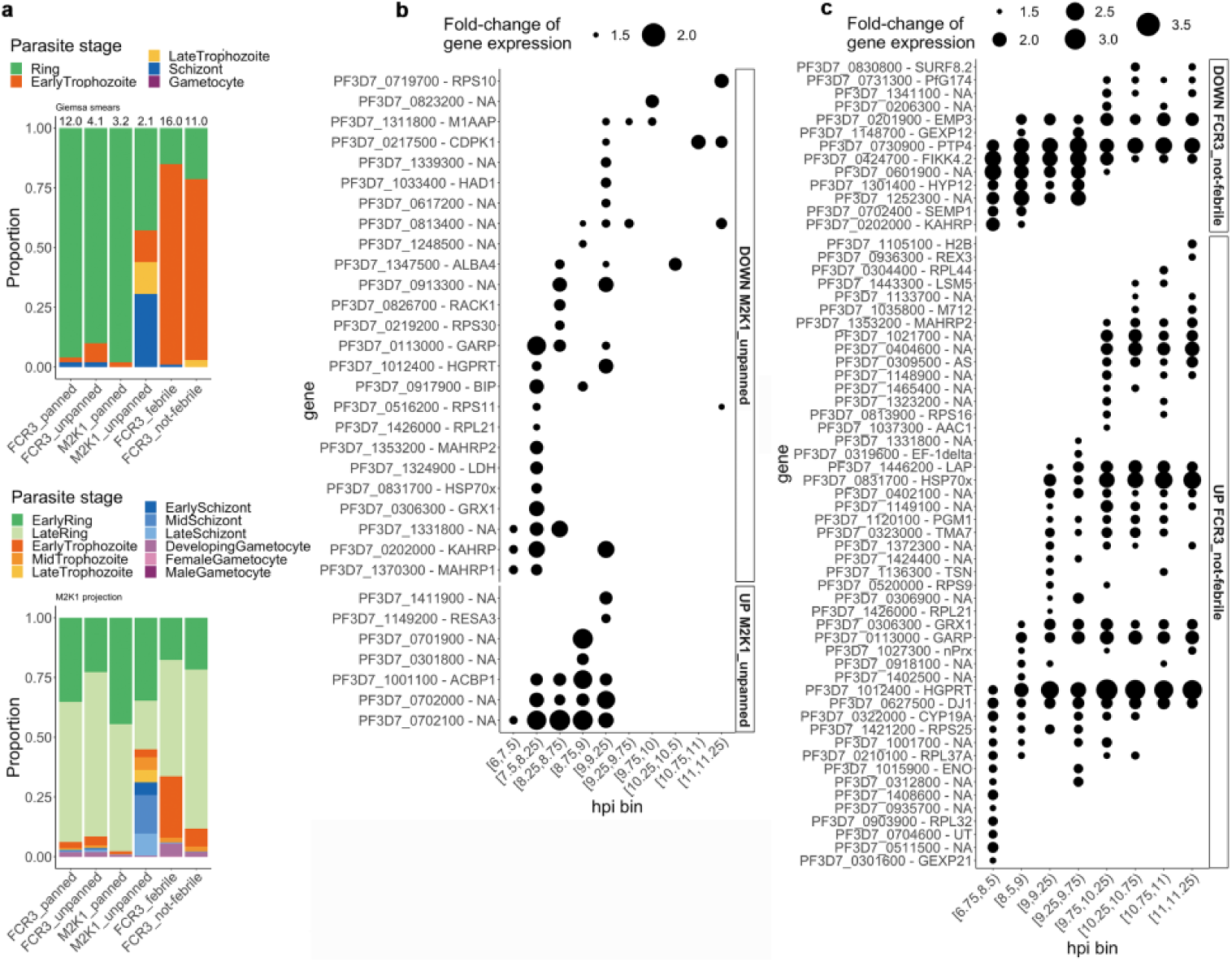
Simulating dry and wet conditions experimentally. **a,** Proportion of parasite stages estimated for each sample by Giemsa-stained thin smears (top) or determined for each iRBC by projection to the M2K1 atlas (bottom). **b,** DE genes down-regulated (top) and up-regulated (bottom) in unpanned vs panned M2K1 iRBCs. Sizes of dots indicate the fold-change of gene expression. **c,** DE genes down-regulated (top) and up-regulated (bottom) in not-febrile vs febrile FCR3 iRBCs. Sizes of dots indicate the fold-change of gene expression.

## Discussion

Recent malaria single-cell atlases include a rodent parasite (17), long-cultured human parasite lines (15), and now a recently adapted 2018 strain (Fig. 1). Each approach may have unique advantages, and collectively their combined data may improve cell type resolution and annotation. Projecting *P. falciparum* transcriptomes from dry season asymptomatic and wet season clinical infections onto these atlases consistently showed older parasite forms in dry season samples (Fig. 2), confirming prior bulk RNAseq findings(7, 35). Nevertheless, natural infections aligned most closely with the M2K1 atlas (Fig. 2d), likely due to its recent adaptation and genetic similarity to Malian strains. The Howick atlas’s lower similarity may stem from fewer iRBCs and cross-species comparisons, while the Dogga atlas’s projections may be influenced by strain differences or long-term culture (44, 45), as it excluded uncultured Malian iRBCs from its annotation (15).

The Dogga (15) and the M2K1 atlases used UMAP and PHATE, respectively, for dimensionality reduction. While UMAP better distinguishes cell types, PHATE captures developmental trajectories (27) but struggled to separate gametocytes from asexual stages (Supplementary Fig. 2), and we relied initially on unsupervised clustering to differentiate sexual from asexual stages, guided by stage-marker transcripts (Fig. 1b, Supplementary Fig. 1).

We benchmarked merozoite reinvasion and intra-erythrocytic progression using marker genes (Fig. 1b), converting pseudotime to estimated hpi. Validation via pseudobulk comparisons (29, 30), confirmed strong developmental correlations (Supplementary Fig. 3), enabling the first hpi-based annotation in a malaria scRNA-seq atlas. The M2K1 atlas was less effective than Dogga’s in identifying mature gametocytes in the natural *P. falciparum* infections, and showed lower similarity to uncultured parasites (median: 0.78, IQR: 0.77-0.79 vs 0.83, IQR: 0.82-0.83), underscoring the need for broader stage coverage and larger cell numbers. While both atlases captured developing gametocytes, projections differed: Dogga aligned with later stages, whereas M2K1 better identified early gametocytes, consistent with van Biljon et al. (31).

Previous bulk RNA-seq studies comparing asymptomatic and clinical parasites (7, 35, 46) were likely limited by the strong impact of stage composition. Our scRNA-seq analysis of stage-matched parasites (6–11.25 hpi) revealed 26 DEGs across 889 genes, effectively controlling for developmental stage-driven expression differences, remaining 14 DEGs that were also detected in the bulk RNAseq (7) (Supplementary Table 4). The overall fewer genes detected are likely due to dropout rates of the single cell approach (47), the narrow window of 6 to 11.25 hpi of our analysis, and sample processing challenges of asymptomatic infections due to lower parasitaemia, requiring extended cell sorting duration (Supplementary Fig. 5).

Early ring stages were predominant in wet-season clinical cases, while late rings and trophozoites dominated dry season persisting infections, precluding differential expression analyses below 6 and above 12 hpi, a gap that could be addressed with larger cell numbers.

Consistent with altered cytoadhesion, DEGs between dry season asymptomatic infections and wet season clinical malaria cases were linked to protein export, host cell remodelling, and cytoadherence (Fig. 3d and e), including key knob-associated genes (*kahrp, fikk4.2, mahrp 1, hsp70x*). While knockout studies suggest these genes enhance adhesion and rigidity (38, 48–50), their upregulation in dry-season infections correlates with prolonged iRBC circulation (Fig. 3) (7). In line with previous reports, *kahrp*, *mahrp1*, and *hsp70x* were elevated in *Pf*M2K1 iRBCs selected for high binding (Fig. 5b). While conditional gene expression studies of these genes have not been reported, one study reported that knob density appeared linked to the *var* variant expressed independently of *kahrp* expression (51), while a different report suggested host endothelial receptor and febrile temperature could select for increased *kahrp* expression and presence of knobs (52). A bulk RNA-seq study comparing *P. falciparum* of non-severe vs severe malaria cases and statistically normalizing the parasite’s developmental stage found *kahrp* and *hsp70x* upregulated in non-severe malaria (53) in line with our observations. However imperfect normalization may have confounded the results. Similarly, our DEGs could reflect residual stage or quality differences between conditions.

Among the 12 genes downregulated in dry-season *P. falciparum*, several were early transcribed membrane proteins (ETRAMPs), with *etramp14* consistently lower across all timepoints (Fig. 3d). ETRAMPs localize to the parasitophorous vacuole membrane (PVM) (54, 55), and *ETRAMP14* upregulation in severe malaria has been linked to PfEMP1 export (56).

RESA, which protects parasites from febrile stress (57), was also downregulated in asymptomatic infections. Since RESA reduces iRBC deformability and slows microcirculation in vitro (58), its lower expression may contribute to prolonged dry-season circulation. While febrile-temperature experiments with *Pf*FCR3 (a *resa*-disrupted line (59)) were inconclusive, *resa* was downregulated in heat-stressed NF54 iRBCs (43), suggesting its clinical upregulation may reflect selection for altered microcirculatory properties rather than direct fever response. Febrile conditions in *Pf*FCR3 cultures induced 61 DEGs between 6.75 and 11.25 hpi, including downregulated *garp*, *hsp70x*, and *lap*—consistent with wet-season clinical cases—implying fever-driven suppression. Conversely, *kahrp* and *fikk4.2* were upregulated by heat, suggesting their dry-season induction is fever-independent.

To determine if single-cell expression differences affected KAHRP and MAHRP1 proteins in iRBCs between seasons, we compared confocal microscopy images of freshly collected iRBCs of dry-season asymptomatic infections and wet-season clinical cases. Dry-season iRBCs had larger nuclei, indicating more developed parasites, along with more KAHRP-positive cells and stronger MAHRP1 staining (Fig. 4). KAHRP localized at the RBC membrane, showing dry-season iRBCs circulate despite high KAHRP levels.

We built the first single-cell atlas of a recently isolated *P. falciparum* strain and used it to confirm that dry season infections harbour more developed parasites in circulation, and to compare parasite gene expression between similar developmental stages in dry season asymptomatic infections and wet season clinical malaria cases. We found altered expression in cytoadherence, protein export, and host remodelling genes in persistent parasites. Dry-season iRBCs showed elevated KAHRP and MAHRP1 levels despite similar nuclear size to clinical cases. Our data also suggest that fever during clinical malaria influences *P. falciparum*’s transcriptional profile, indicating adaptations in dry-season persistence.

## Acknowledgments

We thank the residents of Kalifabougou and Torodo, Mali, for their participation in the studies. This work was supported by the Lise Meitner Excellence Programme of the Max Planck Society, and the Division of Intramural Research, National Institute of Allergy and Infectious Diseases, National Institutes of Health. Katrin Lehmann assisted with sequencing of single-cell libraries. Christian Goosmann and Volker Brinkman at the microscopy facility of the MPIIB supported confocal imaging. We thank Michael Lanzer and Hans-Peter Beck for kindly providing the anti-KAHRP and anti-MARHP antibodies, respectively. We acknowledge the use of PlasmoDB (https://plasmodb.org) throughout this study and thank its team for maintaining this valuable resource.”

## Data availability

Raw sequencing data, analysis and visualisation scripts, and gene expression matrices and metadata are available at https://edmond.mpg.de/ under the identifier doi:10.17617/3.RX6VDG. The M2K1 atlas is uploaded to paraCell (60) for interactive analysis and visualisation (https://cellatlas-cxg.mvls.gla.ac.uk/Plasmodiumfalciparum_M2K1_Clinical_isolate_cellatlas/)

## Methods

### PfM2K1 and PfFCR3 in vitro culturing

*P. falciparum* cultures were maintained in fresh human O^Rh+^ erythrocytes at 3 - 5 % haematocrit in complete RPMI medium; RPMI 1640 with L-glutamine and HEPES, 0.24 % Sodium Bicarbonate, 100 μM Hypoxanthine and 25 mg/L gentamycin, supplemented with 0.25 % Albumax II, and 10 % heat-inactivated human O^Rh+^ plasma, at 37°C in a gas mixture of 5 % O_2_, 5 % CO_2_, and 90 % N_2_. To build the M2K1 atlas, we included 13 in vitro cultures of M2K1 applied with different synchronization and enrichment schedules, including flow cytometric sorting, MACS columns, sorbitol synchronization, merozoite isolation, and gametocyte induction as depicted in Supplementary Table 1.

### Synchronization of PfM2K1 cultures

Parasites cultures were synchronized by sorbitol lysis of mature iRBCs stages as previously described (61) or by magnetic activated cell sorting (MACS) enriching for hemozoin-containing mature iRBCs stages as previously described (62). In brief, the sorbitol synchronization was performed using 9 – 10 times the pellet volume of 5 % D-sorbitol, incubated 8 min at 37°C, followed by two washes of 9 – 10 times the pellet volume of incomplete RPMI medium; the MACS synchronization was performed by resuspension of the RBC pellet in incomplete RPMI medium to 5 – 15 % haematocrit, then loaded to LS columns prefilled with 1 – 2 mL incomplete RPMI medium and held in a magnetic support, followed by washing with 3 – 5 mL incomplete RPMI medium, then elution with 4 – 5 mL incomplete RPMI medium and the column off the magnetic support.

### Merozoite isolation of PfM2K1 cultures

Parasite cultures were synchronized with sorbitol, and merozoites were isolated as previously described (63) with modifications. In brief, parasites at early schizont stage were incubated with 10 μM E64 (64) in the culture medium for 10 hours to block egress. The iRBCs were pelleted (1,900 rcf, 5 min), resuspended in 2 mL incomplete RPMI medium, and pushed through a 1.2 μm syringe filter to mechanically rupture the schizonts. The filtrate was passed directly onto fresh RBCs in incomplete RPMI medium supplemented with 25 % heat-inactivated human plasma and incubated for 30 min at normal conditions to allow reinvasion, then pelleted (300 rcf, 3 min) and cultured in complete RPMI.

### Gametocyte induction of PfM2K1 cultures

Parasite cultures were cultured to high parasitaemia (>5 %, day 0), and at day 5 – 7 incubated with 30 U/mL heparin-sodium for 1.5 cycles to inhibit reinvasion and thereby asexual growth. Gametocytes were used on day 10 – 14.

### Samples of natural P. falciparum infections

The samples of *P. falciparum* infections for single-cell transcriptional analysis were obtained between 2018 and 2020 in a cohort study in Kalifabougou, Mali, and the samples of *P. falciparum* infections for single-cell protein analysis by microscopy were obtained in 2023 in a cohort study in Torodo, Mali. The Kalifabougou study site and cohort design were previously described (33) and included ∼600 individuals from 3 months to 40 years of age. Written informed consent was obtained from all participants or the parents/guardians of the participating children. The Ethics Committee of Heidelberg University Hospital, the Faculty of Medicine, Pharmacy and Odontostomatology (FMPOS) at the University of Bamako, and the National Institute of Allergy and Infectious disease of the National Institutes of Health Institutional Review Board approved this study. The study is registered at ClinicalTrials.org (identifier NCT01322581).

The Torodo cohort included 250 individuals in the age of 3 months to 16 years. Written informed consent was obtained from all participants or the parents/guardians of participating children. The Ethics Committee of Charité (EA2/264/21), and the Faculty of Medicine, Pharmacy and Odontostomatology (FMPOS) at the University of Bamako (N°2022/20/CE/USTTB/24.01.22) approved this study.

Blood samples were collected from symptomatic individuals presenting with their first febrile malaria episodes, and asymptomatic RDT^+^ individuals. From each individual a thick smear, dried blood spots and venous blood (4 or 8 mL depending on if donor age was below or above 4 years old) drawn into sodium citrate-containing cell preparation tubes were prepared. In the venous blood samples, PBMCs, plasma and RBC pellet were separated by centrifugation. The RBC pellet was washed twice with PBS and used freshly or cryopreserved at −80°C mixed with 1.5x the pellet volume of heat-inactivate AB^Rh+^ serum and 2.5x the pellet volume of glycerolyte.

### Thawing of glycerolyte-frozen iRBCs

Glycerolyte-frozen iRBCs of the Kalifabougou cohort samples were rapidly thawed at 37°C, transferred from the cryo tube to a falcon tube, and pelleted (300 rcf, 3 min). 0.1x the pellet volume of 12 % NaCl solution was dropwise added while gently swirling the tube. After 5 min incubation at RT, 10x the pellet of 1.6 % NaCl solution was added dropwise while gently swirling the tube. The iRBCs were pelleted (300 rcf, 3 min) and washed with 10x the pellet volume of complete RPMI medium.

### Flow cytometric enrichment of iRBCs

20 - 50 μL of RBC pellet was incubated in 1 mL of 5x SYBR Green II, 0.5x APC anti-human CD71 antibody (ThermoFisher, cat# 17-0719-42), and 0.25x APC anti-human Lineage Cocktail (BioLegend, cat# 348803) in PBS for 30 min at 37°C, followed by three washes with 2 mL PBS. The cells were sorted on a BD FACSAria III instrument with 70 μm nozzle, into 20 μL of PBS. SYBR^+^ and APC^-^ cells were selected for sorting. 5 – 10 μL were used for post-sort assessment. The flow cytometric data was acquired using BD FACSDiva (v8.0+) and analyzed using BD FlowJo (v10.0+).

### Single-cell capture and preparation of 10x chromium libraries

Samples were partitioned to single-cells according to the manufacturer’s instructions, on 10x Genomics’ Chromium Single Cell Controller using the Chromium Next GEM Chip G Single Cell Kit (10x Genomics, PN-1000120) and Chromium Next GEM Single Cell 3’ Reagent Kit v3.1 (10x Genomics, PN-1000121), and further prepared for Illumina sequencing according to the manufacturer’s instructions using dual indexes (Dual Index Kit TT Set A, 10x Genomics, PN-1000215). The cDNA was amplified in 14-20 cycles and libraries were amplified in 14-16 cycles. Quality assessment of cDNA and libraries was done using the Bioanalyzer 2100 DNA High Sensitivity assay and the Qubit4 High Sensitivity dsDNA assay.

### Globin depletion and Sequencing of 10x chromium libraries

Multiplexed libraries of natural infection samples were depleted for globin inserts with the CRISPRclean Single RNA Boost Kit (Jumpcode Genomics, KIT1018) using guides from the CRISPRclean Globin Depletion Kit (Jumpcode Genomics, KIT1024) as instructed by the manufacturer, using 5 – 10 μL globin guide RNA per reaction, and 8 – 10 amplification cycles post depletion. Libraries were quantified using the Bioanalyzer 2100 DNA High Sensitivity assay and the Qubit4 High Sensitivity dsDNA assay, and sequenced on Illumina NextSeq2000 with P2 or P3 Reagent Kits and the read protocol 26, 10, 10, 90 or 28, 10, 10, 90.

### Mapping of sequencing reads

The *mkref* function of CellRanger (v5.0.1) (65) was used to create a custom reference including the reference genomes of *P. falciparum* 3D7 (PlasmoDB, v60) (66) and Human (UCSC (67), GRCh38/hg38). Gene annotations were obtained from PlasmoDB (v60) and GENCODE (v34) (68). The *count* function of CellRanger was used to map the sequencing reads to the custom reference and create a matrix of UMIs per cell barcode. The UMIs of Human genes were excluded for downstream QC and analysis, which was performed using R Statistical Software (v4.1.0) (69).

### Identification and quality-filtering of iRBCs

Cell barcodes representing iRBCs were determined by the number of genes detected for each barcode; 200 genes or more in samples of in vitro cultures, and 100 genes or more in samples of natural infections. Cell barcodes of iRBCs were further quality-filtered by retaining barcodes less than 20,000 UMIs and less than 5 % mitochondrial UMIs. Additionally, for the M2K1 atlas samples, iRBCs were only retained for samples with more than 200 quality-filtered iRBCs, which excluded two of thirteen samples (Supplementary Table S1). UMIs of rRNAs were excluded for downstream analysis. rRNA ids: PF3D7_0112300, PF3D7_0112500, PF3D7_0112700, PF3D7_0531600, PF3D7_0531800, PF3D7_0532000, PF3D7_0725600, PF3D7_0725800, PF3D7_0726000, PF3D7_0830000, PF3D7_0830200, PF3D7_1148600, PF3D7_1371000, PF3D7_1371200, PF3D7_1371300, PF3D7_1418500, PF3D7_1418600, PF3D7_1418700, PF3D7_API04900, and PF3D7_API05700.

### Integration and cluster-specific quality-filtering of the M2K1 atlas iRBCs

The gene expression data of iRBCs in each sample was normalized using the *quickCluster* method in scran (v1.20.1) (70), and was integrated across samples using Stacas (v2.0.1) (71) with 2,000 anchors. Seurat (v4.3) (72, 73) was used for principal component analysis and clustering of integrated data. Six clusters were identified using the SLM algorithm (v1.3.0) (74) at a resolution of 0.1 (modularity > 0.95) based on a shared nearest neighbour graph computed from principal component 1 – 30 with 100 trees and k.param = 30. The iRBCs of one cluster was excluded due to overall few UMIs per cell, and the iRBCs of the remaining five clusters were filtered based on the standard deviation (sd) from the average number of genes and UMIs per cell within each cluster, retaining iRBCs with more genes than 0.7 sd below the average number of genes, and less UMIs than 2 sd above the average number of UMIs. The normalization, integration, principal component analysis, and clustering were recalculated for the retained iRBCs as described above, but using a cluster-resolution of 0.5 identifying thirteen clusters. The integrated gene expression data was denoised using *magic* in Rmagic (v2.0.3) (18) with t=’auto’.

### Dimensionality reduction and trajectory/pseudotime inference of the M2K1 atlas iRBCs

PHATE dimensionality reduction was performed using *phate* in phateR (v1.0.7) (27) with mds.method=’nonmetric’, ndim=3, and t=100, and nine KMeans (75) clusters were identified with centers=9 and nstart=5. Trajectories were inferred using *slingshot* (v2.0.0) (28) with the input of the PHATE embedding and the nine KMeans clusters. The analysis done with all iRBCs together, but eventually done separately for iRBCs of sexual and asexual SLM clusters determined by stage marker genes.

### Annotation of the M2K1 atlas iRBCs

The M2K1 atlas iRBCs were annotated by the separate trajectory and pseudotime inference of asexual and sexual iRBCs. The pseudotime of asexual iRBCs was scaled to the 48h IDC by dividing the maximum pseudotime value and multiplying with 47.999 to obtain annotation by hpi. The asexual iRBCs were grouped by hpi into discrete parasite developmental stages: early ring (0–10 hpi), late ring (10–15 hpi), early trophozoite (15–21 hpi), mid trophozoite (21–26 hpi), late trophozoite (26–32 hpi), early schizont (32–38 hpi), mid schizont (38–44 hpi), and late schizont (44–48 hpi). The sexual iRBCs showed two trajectories and were annotated by iRBCs which were shared between trajectories (developing gametocyte) and iRBCs which were unique to each trajectory (female gametocyte or male gametocyte).

### Comparison of the M2K1 atlas iRBCs to bulk transcriptome datasets

The asexual iRBCs were grouped by hpi into 48 bins of 1h intervals. The gene expression of iRBCs in each bin were summed to a pseudo-bulk sample and compared to microarray 48h IDC time-course data (29) using an MLE model (30). The sexual iRBCs were grouped by annotated stage and further sub-grouped by pseudotime of equal intervals into six bins of developing gametocytes (DG1 – DG6), two bins of female gametocytes (FG1 and FG2), and one bin of male gametocytes (MG). Expression of iRBCs in each bin were summed to a pseudo-bulk sample and compared to RNA-seq 16-day time-course data of sexual commitment and gametocyte development (31) by Spearman’s correlation.

### Comparison of the M2K1 atlas to existing single-cell atlases

The M2K1 atlas was compared to the malaria cell atlases of *P. berghei* (17) and of *P. falciparum* (15) using the *scmapCell* function of scmap (v1.14.0) (32). The gene expression data was log1p-normalized, and the iRBCs of each atlas were indexed using 500 genes. The *P. berghei* genes were converted to the orthologous *P. falciparum* genes according to the ortholog table provided in the supplementary material (17). The queried iRBC was assigned the metadata of the top projected index cell, and the cosine similarity was recorded. The cosine similarity obtained by projection to different atlases was compared by Wilcoxon signed rank test.

### Comparison of existing single-cell atlases to bulk transcriptome datasets

Asexual iRBCs of the Howick atlas were grouped by the reported pseudotime into 48 bins, and the gene expression of iRBCs in each bin were summed to a pseudo-bulk sample and compared to microarray 48h IDC time-course data (29) using an MLE model (30).

The Dogga atlas had no reported pseudotime and was not compared.

### Screening for additional Plasmodium spp. in natural P. falciparum infections

The *mkref* function of CellRanger (v5.0.1) (65) was used to create a custom reference including the reference genomes of *P. falciparum* 3D7, *P. vivax* P01, *P. ovale curtisi* GH01, *P. malariae* UG01, and *P. knowlesi* H (PlasmoDB, v60) (66), and additionally Human (UCSC (67), GRCh38/hg38). Gene annotations were obtained from PlasmoDB (v60) and GENCODE (v34) (68). The *count* function of CellRanger was used to map sequencing reads to the custom reference and create a raw gene count matrix of UMIs per cell barcode. The UMIs of Human genes were excluded, and the remaining UMIs were used to quantify the proportion of each *Plasmodium sp.* in each cell. Only cell barcodes showing at least 95 % *P. falciparum* UMIs were used for downstream analysis.

### Stage-determination of P. falciparum-iRBCs of natural infections and of in vitro cultures

Parasite developmental stages of *P. falciparum*-iRBCs were determined by projection to the M2K1 atlas using the *scmapCell* function of scmap (v1.14.0) (32). The gene expression data was log1p-normalized, and the iRBCs of the M2K1 atlas were indexed using 500 genes. Each queried iRBC was assigned the metadata of the top projected index cell, and the cosine similarity was recorded.

### Stage-matching of P. falciparum-iRBCs and differential gene expression analysis

For each comparison, the iRBCs were initially binned by hpi at intervals of 15 min. Then, neighbouring bins with less than 100 stage-matched iRBCs were merged. Bins with at least 100 stage-matched iRBCs and a maximum hpi interval of 1.5h were considered for differential gene expression analysis.

The gene expression of the iRBCs in each comparison was normalized using the *quickCluster* method in scran (v1.20.0) (70). Within each hpi bin of iRBCs, differential gene expression between sample groups was tested using the MAST method (v1.18.0) (76) in Seurat (v4.3) (72, 73) for genes expressed in at least 10 % of iRBCs in either sample group. DE genes were determined by BH-adjusted P value < 0.05, fold-change > 1.5, and gene expression in at least 10 % of iRBCs in both sample groups within at least one of the tested hpi bins.

### Gene ontology enrichment analysis

Gene ontology (GO) terms were tested for enrichment within each hpi bin using the *gsea* function of clusterProfiler (v4.0.5) (77) using the GO database of *P. falciparum* 3D7 (PlasmoDB, v60) (66). Genes were ranked by average log2 fold-change. Only GO terms with at least five genes were considered.

GO terms were significantly enriched by BH-adjusted P value < 0.05.

### Immunofluorescent assay of thin blood smears

PBS-washed RBC pellets of the Torodo cohort samples, were used fresh to prepare thin blood smears, which were incubated in −20°C 100 % methanol for 7 min to fixate and permeabilize the cells. The smears were washed briefly in dH_2_O (twice), air-dried and stored in airtight containers with desiccant at −20°C until further use. The smears were first incubated in blocking buffer (3 % BSA and 0.05 % Tween20 in PBS) for 2 hours at 4°C; then stained with primary antibodies (1:500 rabbit anti-KAHRP and 1:500 mouse anti-MAHRP1 in blocking buffer) O/N at 4°C followed by three washes in blocking buffer for 5 min at 4°C; then stained with secondary antibodies (1:500 Alexa555-goat anti-rabbit (ThermoFisher, cat# A-21428) and 1:500 Alexa647-goat anti-mouse (ThermoFisher, cat# A-21235) in blocking buffer) for 1 hour at 4°C followed by three washes in blocking buffer for 5 min at 4°C; then stained with a conjugated primary antibody (1:500 atto488-mouse anti-EXP2 in blocking buffer) O/N at 4°C followed by one wash in blocking buffer and two washes in PBS for 5 min each at 4°C; then counterstained with Hoechst (1:1000) for 5 min at 4°C followed by two washes in PBS for 5 min at 4°C; then mounted (AquaPoly Mount) O/N at room temperature. The smears were read on confocal Microscopes Leica SP8 and DM6000B-CS. The area of the parasite’s nucleus was measured using Qupath (v0.5.1) with ‘Watershed Cell detection’ function on Hoechst stain, pixel size 0.1, background radius 1.0, median radius 0.0 and the sigma of 0.4 microns. Minimum area was defined as 0.5 and maximum area as 10.0 microns. Intensity threshold value was 40.0. A script was defined and ran for the batch of images resulting in 330 nuclei measurements that were manually curated to verify that all nuclei were inside parasitophorous vacuoles, resulting in a total of 246 measurements (186 measurements for MAL and 60 measurements for endDry) for 125 images (MAL23T211: 14 images, MAL23T197: 18, MAL23T062: 17, MAL23T133: 17, MAL23T142: 2, endDry23T113: 7 images, endDry23T077: 11, endDry23T147: 7, endDry23T225: 9, endDry23T226: 18, endDry23T142: 5). KAHRP positivity was manually defined, and KAHRP radial profile was measured in all 246 cells using the radial profile extended plugin (http://questpharma.u-strasbg.fr/html/radial-profile-ext.html)using imageJ.

MAHRP1+ cells were first defined manually in imageJ using the channels tools, on the Alexa647 channel. We defined 89 MAHRP1+ cells in a total of 74 images and a script was made in image J to measure the total area and the mean intensity of MAHRP1 in MAHRP1+ cells. Images were displayed in Gray Scale, then blurred with the ‘Gaussian Blur’. Sigma was defined in 2 stacks and we set the auto threshold ‘Max entropy’. Minimum size of MAHRP1+ structures was defined as 0.1 and the maximum as infinite. We ran the script for the 74 images that contained MAHRP1+ cells and acquired measurements of count of MARHP1+, size in micrometers of each MAHRP1 dot, their average size, MAHRP1s total area, their mean intensity separately and all together.

### Panning of PfM2K1 and PfFCR3 iRBCs

Parasite cultures were panned on human dermal microvascular endothelial cells (HDMECs) to enrich for adherent iRBCs. HDMECs (Promocell, C-12210) were thawed and cultured according to the manufacturer’s instructions in MV2 medium and in pre-coated flasks with 0.3 mL/cm^2^ MV2. A near-confluent culture of HDMECs was prepared by stimulating with 10 ng/mL TNFalpha in MV2 medium without hydrocortisone for 16 – 24 hours at 37°C and 5 % CO_2_, then washed with HBSS. iRBCs were prepared by mixing the pellet with 1.4x the pellet volume of complete RPMI medium and 2.4x the pellet volume of gelafundin (Praxisdienst), followed by incubation at 37°C for 15 min. The pellet of iRBCs was washed twice with incomplete RPMI medium and resuspended in 0.5 % BSA supplemented RPMI 1640 medium with L-glutamine and HEPES, pH 6.8, then added to the HDMECs and incubated 30 – 45 min at 37°C and 5 % CO_2_ to allow binding, with gentle rotation of the flask every 15 min. Then, the suspension of unbound cells was removed, and the HDMECs with bound iRBCs were washed five times in 2 % FCS/PBS. Complete RPMI medium with 5 % fresh RBCs were added to the HDMECs and incubated O/N at 37°C and a gas mixture of 5 % O_2_, 5 % CO_2_, and 90 % N_2_ to allow reinvasion. The suspension of freshly iRBCs was removed from the HDMECs and moved to normal culture.

### Simulation of febrile growth of PfFCR3 cultures

Malarial fever was simulated as previously described (43) with modifications. In brief, the iRBCs were panned and directly subjected to growth at 41°C for 8h, followed by growth at 37°C for the rest of the cycle. In the beginning of the following two cycles, upon reappearance of ring stage iRBCs, the culture was again subjected to 41°C for 8 hours. The culture of panned iRBCs was initially at ∼1 % parasitaemia and 4 % haematocrit, and was split in two for a control culture that was continuously incubated at 37°C. All incubations happened in a candle jar, and the cultures were monitored for parasitaemia and composition of parasite developmental stages by Giemsa-stained thin smears.

---

**Supplementary Figure 1.**
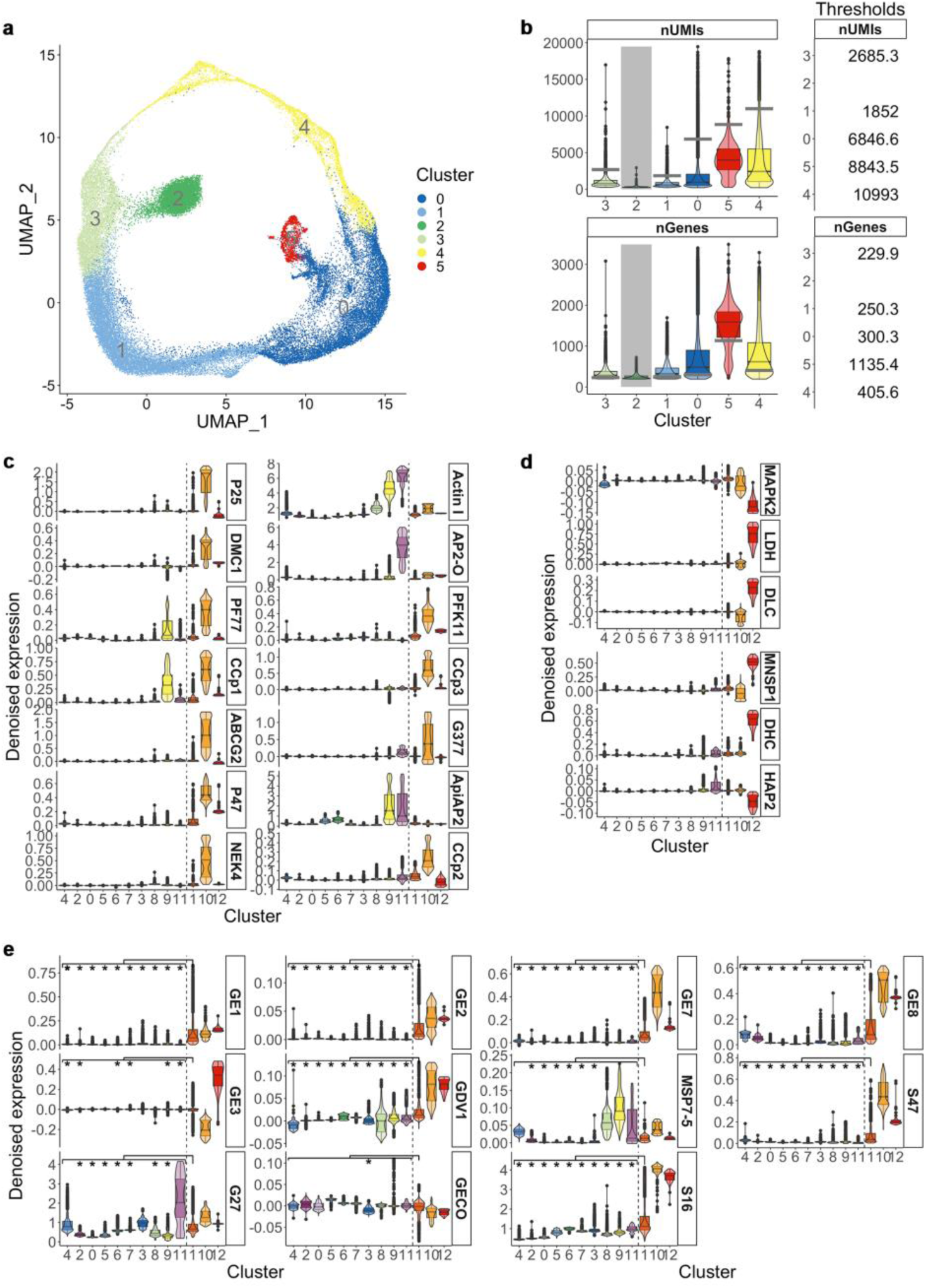
Cluster-specific quality-filtering and marker genes of the M2K1 atlas. **a,** UMAP of 44,813 single cells coloured by six SLM clusters. **b,** Cluster-specific quality thresholds of UMIs (upper limit) and genes (lower limit) per cell indicated by grey bar. Grey shading indicates one excluded cluster. **c,** Female gametocyte and **d,** male gametocyte marker genes. **e,** Early gametocyte development marker. *P < 0.01, pairwise one-sided Wilcoxon rank sum test).

**Supplementary Figure 2.**
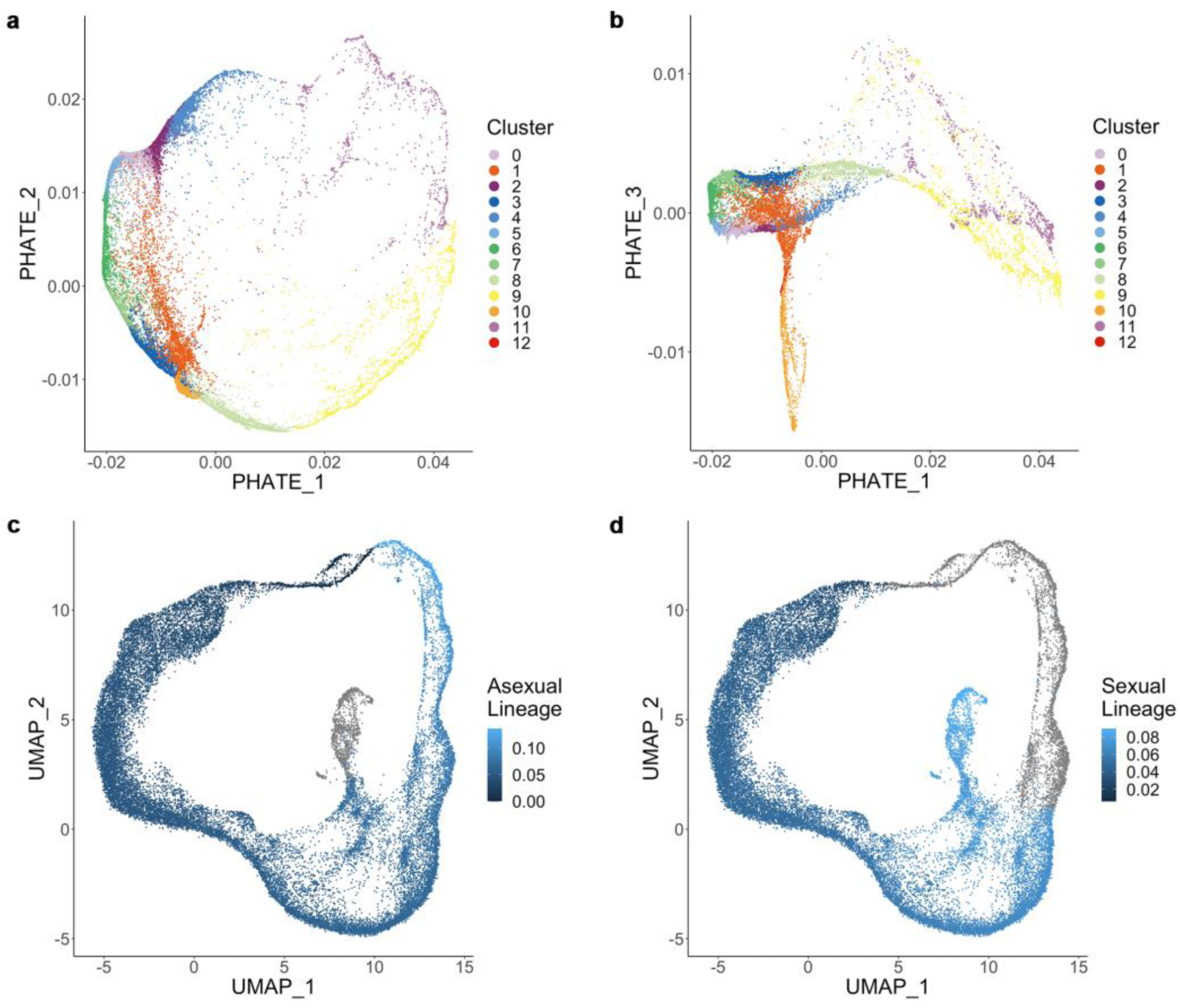
**a and b,** PHATE of 26,477 single cells coloured by thirteen SLM clusters as in Figure 1A, and **c and d,** UMAP coloured by two inferred trajectories/lineages.

**Supplementary Figure 3.**
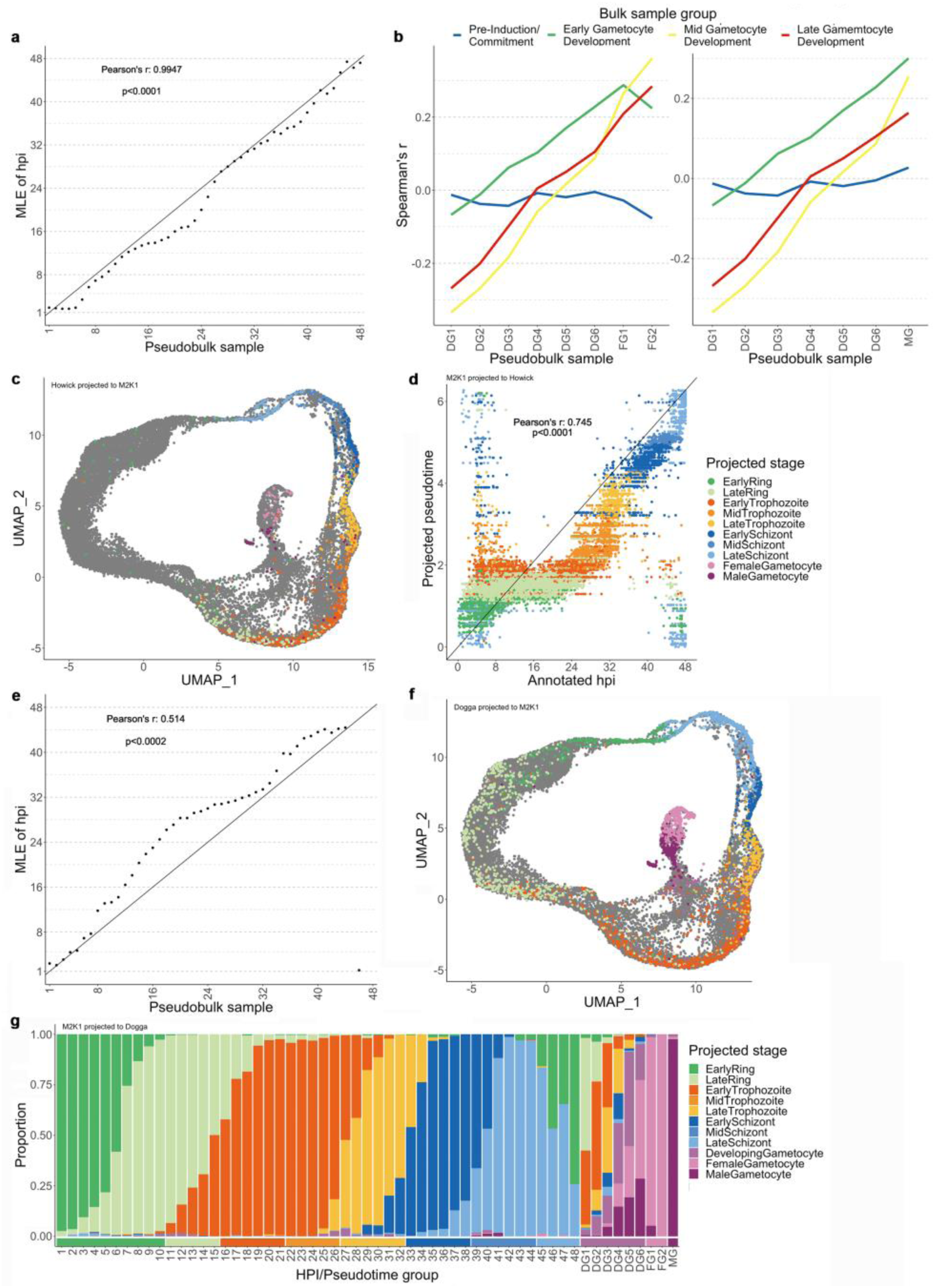
**a,** Correlation of 22,282 asexual cells pseudo-bulked in 48 hpi bins, and the maximum likelihood estimate (MLE) of hpi based on bulk microarray samples of the IDC (Pearson’s r: 0.9947, P < 0.0001). **b,** 4,165 sexual cells pseudo-bulked within pseudotime bins; six bins for developing gametocytes (DG1 – DG6), two bins for female gametocytes (FG1 and FG2, left), and one bin for male gametocytes (MG, right). Correlated to bulk samples of sexual commitment and gametocyte development (Spearman’s r). **c,** UMAP of the projection of the existing *P. berghei* atlas (n = 4,884) to the M2K1 atlas coloured by the annotation in the *P. berghei* atlas. **d,** Correlation of the annotated hpi of 26,447 cells in the M2K1, and the pseudotime according to the projection to the *P. berghei* atlas (Pearson’s r: 0.745, P < 0.0001). **e,** Correlation of the existing *P. berghei* atlas’ 4763 asexual cells pseudo-bulked in 48 hpi bins based on reported pseudotime, and the maximum likelihood estimate (MLE) of hpi based on bulk microarray samples of the IDC (Pearson’s r: 0.514, P < 0.0002). **f,** UMAP of the projection of the existing *P. falciparum* atlas (n = 45,691) to the M2K1 atlas coloured by the annotation in the existing *P. falciparum* atlas. **g,** Composition of annotated stages for the M2K1 atlas’ 26,447 cells in hpi/pseudotime bins according to the projection to the existing *P. falciparum* atlas. Bottom horizontal bar indicates the M2K1 atlas annotation.

**Supplementary Figure 4.**
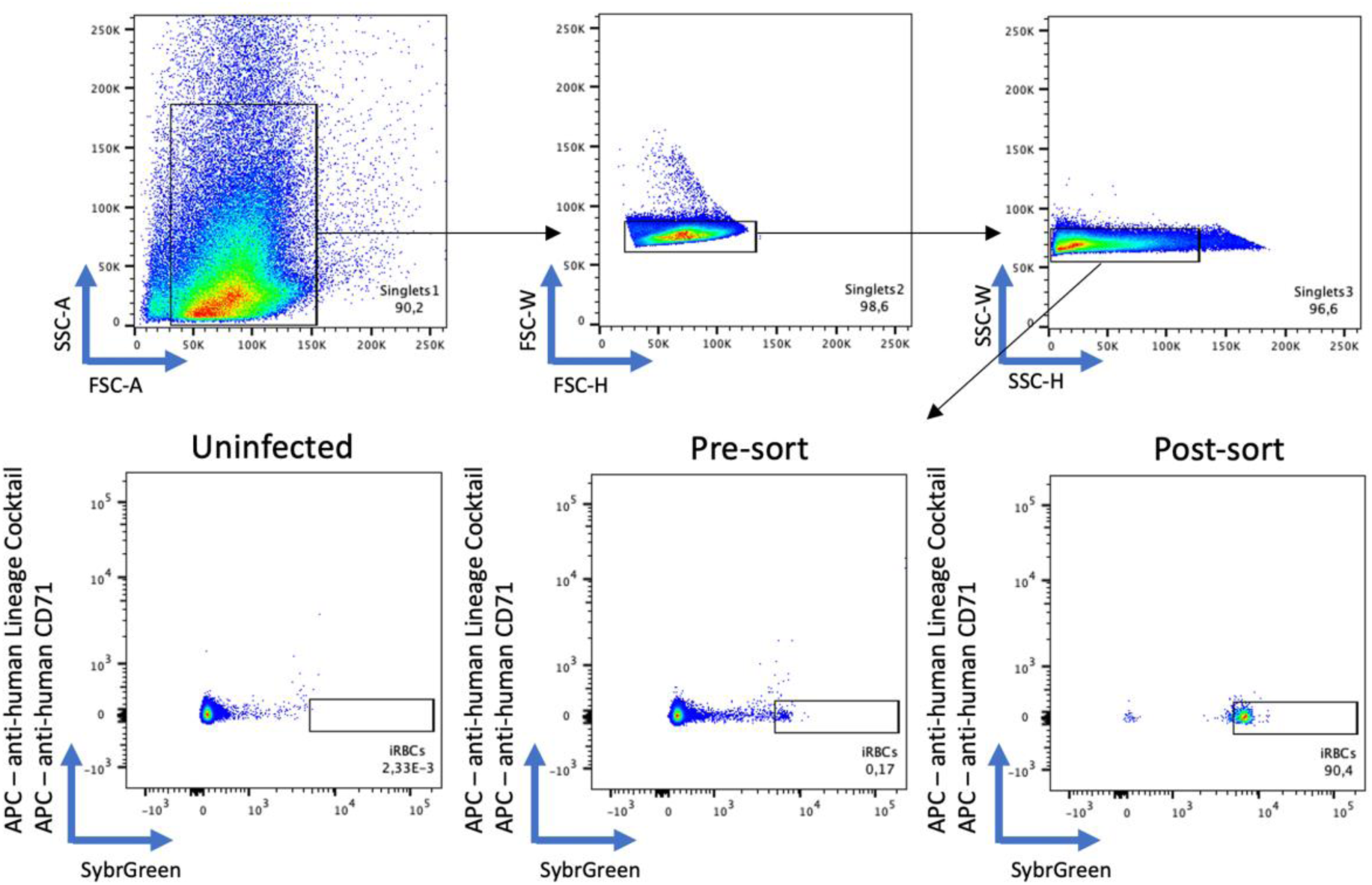
Gating strategy used to flow sort iRBCs. Single event RBCs were selected based on FCS and SSC parameters (top), and iRBCs defined as cells positive for nuclear staining (SYBR^+^) and negative for lineage marker and CD71 (APC^−^) (bottom). Example of uninfected and infected pre- and post-sort are shown.

**Supplementary Figure 5.**
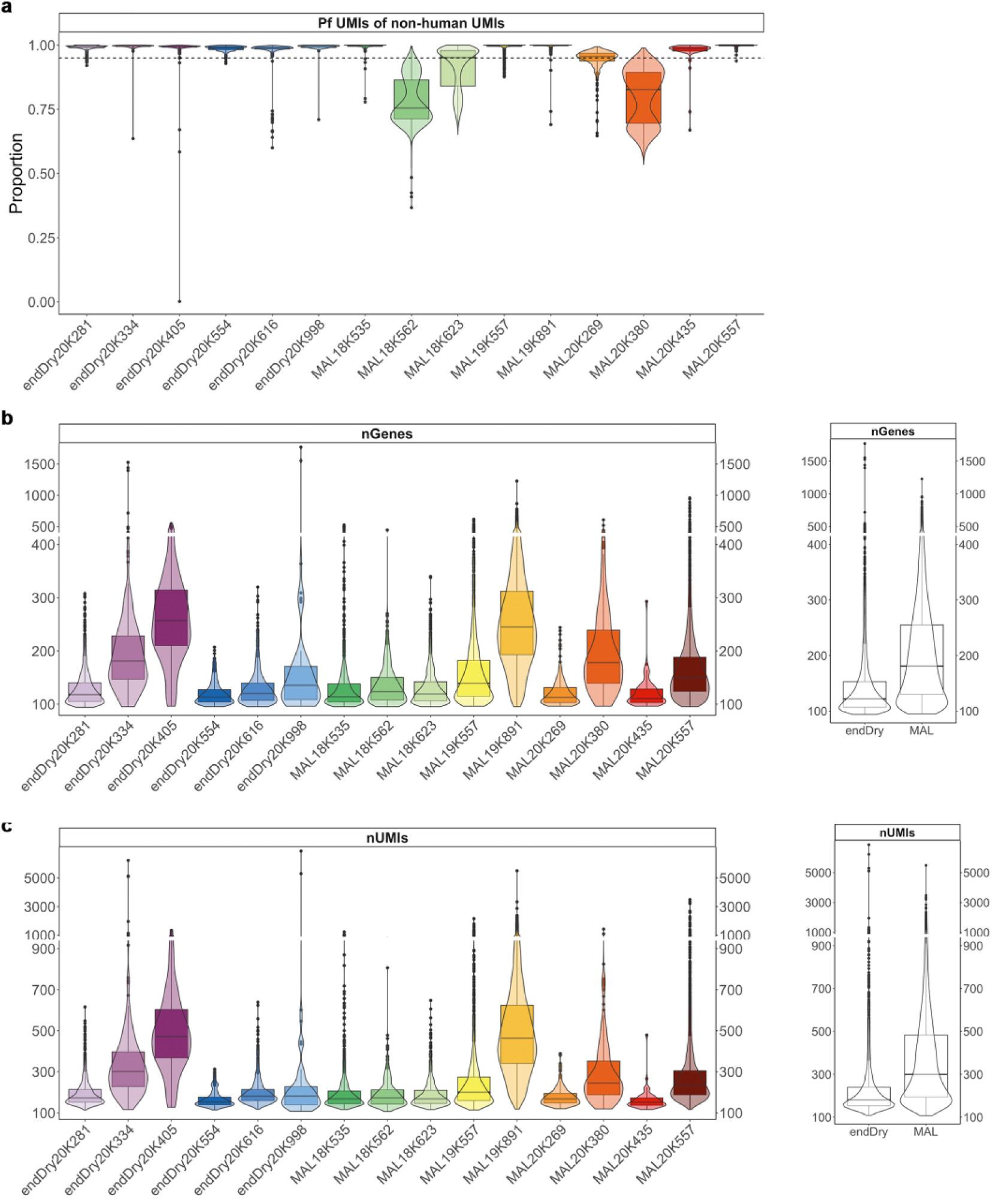
**a,** Proportion of *P. falciparum* UMIs according to the mapping to the five human *Plasmodium spp.* in 3723 single-cells from six asymptomatic infections at the transition from dry to wet season (endDry) and 46221 single-cells from nine clinical malaria cases in the wet season (MAL). Dashed line indicates 95 % threshold. **b,** UMIs and **c,** genes per cell within 3688 endDry single-cells and 23207 MAL single-cells.

## Notes

https://cellatlas-cxg.mvls.gla.ac.uk/Plasmodiumfalciparum_M2K1_Clinical_isolate_cellatlas/

### Competing Interest Statement

The authors have declared no competing interest.

